# *Host_microbe_mapper* allows to analyse and interpret the expression of dual RNA-seq measurements and reveals potential microbial contaminations in the data

**DOI:** 10.1101/234278

**Authors:** Thomas Nussbaumer

## Abstract

Next-generation sequencing technologies provide a wealth of sequencing data. To handle these data amounts, various tools were developed over the last years for mapping, normalisation and functional analyses. To support researchers in the interpretation of expression measurements originating from dual RNA-seq studies and from host-microbe systems in particular, the computational pipeline *host_microbe_mapper* can be applied to quantify and interpret dual RNA-seq datasets in host-microbe experiments. The pipeline with all the required scripts is stored at the Github repository (https://github.com/nthomasCUBE/hostmicrobemapper).

## Background

Different RNA-seq technologies generate huge amounts of data in a cost-efficient manner. To support researchers during this analysis, various computational methods can process the data such as methods for quantification and functional interpretation. For the mapping of single or paired-end reads, the tools STAR [1], Tophat [2] and *ballgawn* [3] are commonly applied in eukaryotic genomes whereas *bowtie* [4] and *bwa* [5] are commonly used for the mapping of microbial reads.

After reads have been assigned to the genome, various ways to interpret the data exist, but it needs a normalisation step before. Whereas in the *cufflinks’* [6] package, there are options to directly obtain FPKM (Fragments per Kilobase Million) counts that consider the gene length and the paired-end information, for other tools it needs to include additional programs to obtain raw read counts which can be extracted with tools such as *featureCounts* [7] or HT-seq [8]. This allows to normalize the data into TMM (Transcripts Per Million Mapped) or RPKM (Reads Per Kilobase Million) in order to perform *e.g*. the detection of differentially expressed genes or to obtain clusters of co-expressed genes with *WGCNA* [9]. Differentially expressed genes can be computed with tools such as *EdgeR* [10], *Noiseq* [11], and *DESeq2* [12] whereas *cufflinks* and *ballgawn* allow to detect differentially expressed genes as part of their pipelines. Most of the mentioned tools have been included into the current pipeline.

An additional aspect of the mapping of expression can be the removal of ribosomal RNA, which can contribute a major proportion of reads in a sequencing run. Reads of ribosomal origin can be discarded from most experiments with available kits (*e.g*. RiboZero Gold kit [13]) or *in silico* by tools such as *SortMeRNA* [14]. Otherwise, it might be of major interest to detect putatively unexpected contaminations in the data. To analyse contaminations based on 16S rRNA genes, we have also included *SortMeRNA* [14] to classify reads, that belong to eukaryotic or prokaryotic ribosomal genes. To reveal or confirm the availability of certain microbes it might be of interest to reconstruct the entire 16S or 18S ribosomal gene from the transcriptomic data. For this aim, there are tools such as *REAGO* [15] that allows to assemble these reads. Otherwise, for dual RNA-seq experiments, it can be crucial to verify, that the pathogen is the main microbial source in the dataset and host response might not active because of other microbes.

Before the entire read set is used to obtain an insight into the data, it needs preliminary access to the data to grasp a first insight into the dataset and to define most appropriate parameters that can be applied on the entire dataset. This requires much more time and computational resources in the following even so a test set might already give a good overview of the data already. This can be ideally done by randomly selecting reads (*e.g*. 100,000) from the dataset. To ease the usage of mapping dual RNA-seq data, we provide a simple graphical user interface that allows to define the reference genomes and to define whether the full or a subset should be used. Despite a bunch of tools that provide a guideline for mapping dual RNA-seq experiments, the pipeline as described in this study has the advantage that is can be run directly on the data without need for major changes as we demonstrate in this study. Calculations can be also run in parallel by using the SLURM batch system. We demonstrate the pipeline by selecting running the experiments on various published different datasets.

## Material and Methods

### Data access

RNA-seq datasets and respective genomic sequences from human, mouse and the pathogen *Neisseria meningitidis* were obtained from NCBI server and Gene Bank identifiers are summarized in **Supplemental Table 1** and were used to obtain artificial read datasets with the ArtificialFastqGenerator.jar by using the coding sequences of the respective genomes [16]. All datasets represent paired-end sequences with 300 nt inserts. In total, 20 million reads were generated for each of the hosts as well as two million reads from the microbe *Neisseria meningitidis*. The respective demo files are included on the Github page.

### Pipeline construction

The whole pipeline is summarized in a single batch script, labelled as *host_microbe_mapper*, that contains all the relevant commands starting from the pre-processing of the data, the mapping of host and microbial reads and finally the data normalisation. All scripts are stored in the Github repository (https://github.com/nthomasCUBE/hostpathogenmapping). Additional prepared scripts can be used to obtain the differentially expressed genes. Therefore, we integrated scripts to perform differential expression analyses by comprising methods such as *EdgeR*, *NOISeq* and *DESeq2*, however these scripts might need additional information and mapping files between samples and conditions. The whole functionality and mapping was performed on the Leibnitz Rechenzentrum Cluster.

## Results and Discussion

### General description of the pipeline

The entire workflow of the program is given in **Figure 1**. A user can provide reads either in the raw FASTQ format or in the compressed format (in the GZ format) by defining the path to the directory that contains them by running the graphical user interface. Each sample within that directory is serially processed. Before reads are mapped to the respective genomes, we perform a quality assessment by checking the FASTQ sequences with *FastQC* [17], followed by adapter removal with *cutadapt* [18]. Next, reads are cleaned from ribosomal reads by considering the *SortMeRNA* pipeline [14] that reports reads matching to ribosomal genes from both, eukaryotes and prokaryotes and are assembled with REAGO [15]. Next, reads are mapped first to the microbe and remaining reads are then mapped to one or two hosts. This is case, when *e.g*. human DNA material is grafted into a mouse model, thereby three different organisms can be simultaneously mapped. **Supplemental Table 2** lists the amount of reads in total that were mapped for the individual datasets and the reads that remain after adapter removal, pre-processing and mapping. These information are collected inside a project-specific Excel file to provide an intuitive overview of the mapping results and furthermore visualisations are made to depict how many of the reads are uniquely mapped, multiple mapped or remain unmapped (**Figure 2A**). For the mapping of eukaryotic reads, STAR [1] or *Tophat* [2] are used and can be selected by the user. Before, reads are mapped to the microbe by using either *BWA* or *bowtie*. Finally, remaining reads are used and are normalised and then differentially expressed gene are applied (*EdgeR* and/or *DESeq2*). For each individual step in listed in separate script files.

**Figure 1.**
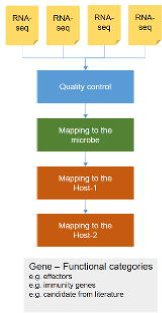
Illustration of the next-generation sequencing (NGS) pipeline covering the steps from the integration of the data to the mapping to the microbe and furthermore the mapping to the reference genomes from both hosts.

**Figure 2.**
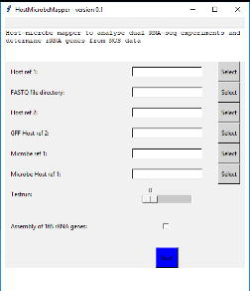
Graphical User Interface for mapping RNA-seq data for the eukaryotic and prokaryotic genomes preparing the mapping files that can be run then on the server.

### Analyses of the dual RNA-seq datasets from published studies

Artificial datasets were generated (see Material and Methods) including dual RNA-seq datasets, that contain a low, median and a high number of microbial reads being present at 1%, 5% or 20%. We used human and mouse expression transcriptomic datasets to simulate these reads from the respective human, mouse genes and pathogen. Then, we applied the *host_microbe_mapper* pipeline and compared known expression counts to the expression, that was from measured from the various mapping tools.

### Possibility to assign the 16S genes from the transcriptome dataset directly

As an additional feature, *host_microbe_mapper* can extract reads from the transcriptome dataset itself to assess which microbes appear in the original raw data, *e.g*. to reveal putative contamination or to detect *e.g*. endophytes, that co-exist with the host. We extracted the classified reads from *SortMeRNA* to extract and then to assemble the entire 16S rRNA genes which can be then compared to reveal existing microbial taxa from the data. To test the pipeline, we have used again reads from the host but now added and simulated the reads from the different microbes. To simulate this, we included microbial datasets from other studies and checked then whether we were able to find them again. Typically use cases for this method would be experiments, that were run in the same batch and thereby might, given misleading barcoding, contain a mixture of reads from different experiments.

